# Scaling contact force parameters across body size, limb count, and number of contact spheres

**DOI:** 10.1101/2025.11.26.690874

**Authors:** Pasha A. van Bijlert

## Abstract

A popular way to model contact interactions in musculoskeletal simulations uses Hertz theory applied to contact spheres, with Hunt Crossley based dissipation. Suitable contact parameters for dynamic simulations will be highly dependent on the morphology, scale, materials, and movement in question. Inappropriate parameter choices can manifest in unpredictable ways during simulations, potentially resulting in misinterpretations or failed simulations.

Here, I demonstrate that both the plane strain modulus and the dissipation parameters are not scale invariant. I derive equations to scale the contact parameters in dimensionless form, which allows accounting for differences in body size, number of legs, contact sphere radius, and number of spheres per foot. As a demonstration of this scaling approach, I scale the contact parameters of a 62 kg human to a 500 kg human, a mouse (0.02 kg), an emu (37.8 kg), a horse (545 kg), and a giraffe (1190 kg), and demonstrate that geometrically and dynamically similar contact behaviour is achieved in all cases.

The scaling approach presented here can be used to scale parameters known to work for one model to a completely different model, which is particularly useful in studies that simulate the effects of allometric scaling. I also provide equations to estimate suitable contact parameters for a model directly, without using a different model as a starting point. The limitations of Hertz Hunt Crossley contact models in biomechanical simulations are discussed. Lastly, I derive dimensionless expressions and scaling guidelines for the smoothed contact force implementation “SmoothSphereHalfSpaceForce” in the popular biomechanical simulator OpenSim.

## 1. Introduction

Dynamic simulations of musculoskeletal models have a wide variety of fundamental and clinical applications in biomechanics (Van Soest et al., 1993; van den Bogert et al., 1989; Sellers et al., 2013; Bianco et al., 2023). It is often required to model contact interactions, for example to acquire ground reaction forces during locomotion (van den Bogert et al., 1989; Sellers et al., 2013; Bianco et al., 2023; Bates et al., 2025) or forces on the teeth during biting (Watson et al., 2014; Bates and Falkingham, 2012). A popular approach to achieve this is using contact spheres (Sherman et al., 2011; Seth et al., 2018; Van Bijlert et al., 2024; Clemente et al., 2024; van Bijlert et al., 2024; Rodrigues Da Silva et al., 2022). When these spheres intersect with other spheres or a halfspace (i.e., a plane), the elastic portion of the forces are computed following Hertz theory of elastic contact (Hertz, 1896; Johnson, 1985; Sherman et al., 2011) ^1^. Dissipation is incorporated as a function of both the elastic force and the intersection speed, using the Hunt Crossley model (Hunt and Crossley, 1975).

Choosing suitable contact parameters for simulations often represents a tradeoff between realism and numerical performance (van den Bogert et al., 1989). High stiffness in the contacts leads to stiff differential equations, which is numerically undesirable because it causes long computation times and may lead to locally-optimal solutions or even failed simulations. Conversely, very compliant contacts tend to be easier to simulate, but may no longer accurately represent physical interactions experienced by the animal, affecting the simulated kinematics. Furthermore, contact forces tend to be modelled as one of the primary sources of energy dissipation in the system (Afschrift et al., 2025), directly influencing the energetics of a movement. Thus, unsuitable contact parameters will have negative effects on simulations, which could be misinterpreted as biologically meaningful results.

Unfortunately, judicious parameter selection is not straightforward when using contact spheres. Often, the composite plane strain modulus *E*^*^ is incorrectly treated and referred to as “stiffness”, which is a derived property, whereas *E*^*^ is a material property (see sec. 4.3). In Hertz contact mechanics, stiffness will also vary non-linearly with the radius of the contact spheres, and their relative deformations. Similarly, the Hunt Crossley dissipation parameter affects the coefficient of restitution in a scale dependent way.

Here, I will derive a scaling equations for contact parameters based on Hertz theory of elastic contact, combined with Hunt Crossley dissipation. The scaling equations result in geometrically and dynamically similar contact sphere interactions at different body sizes, number of spheres per foot, and contact radii, allowing for size-invariant contact behaviour. I will demonstrate the effects of this scaling using simulated drop tests of a variety of animal models of varying body sizes and shapes (Fig. 1). Simulations were performed using the popular simulator OpenSim (Seth et al., 2018; Dembia et al., 2020), which implements Hertz-style contacts (with Hunt Crossley style dissipation (Hunt and Crossley, 1975)) as both “HuntCrossleyForce” and “SmoothSphereHalfSpaceForce”. Additional derivations for size-invariant behaviour of “SmoothSphereHalfSpaceForce” are presented in Appendix C. While this manuscript is primarily concerned with contact interactions in musculoskeletal systems, the underlying scaling principles are general and applicable to any system modeling Hertzian elastic contact with Hunt Crossley inspired dissipation.

**Figure 1.**
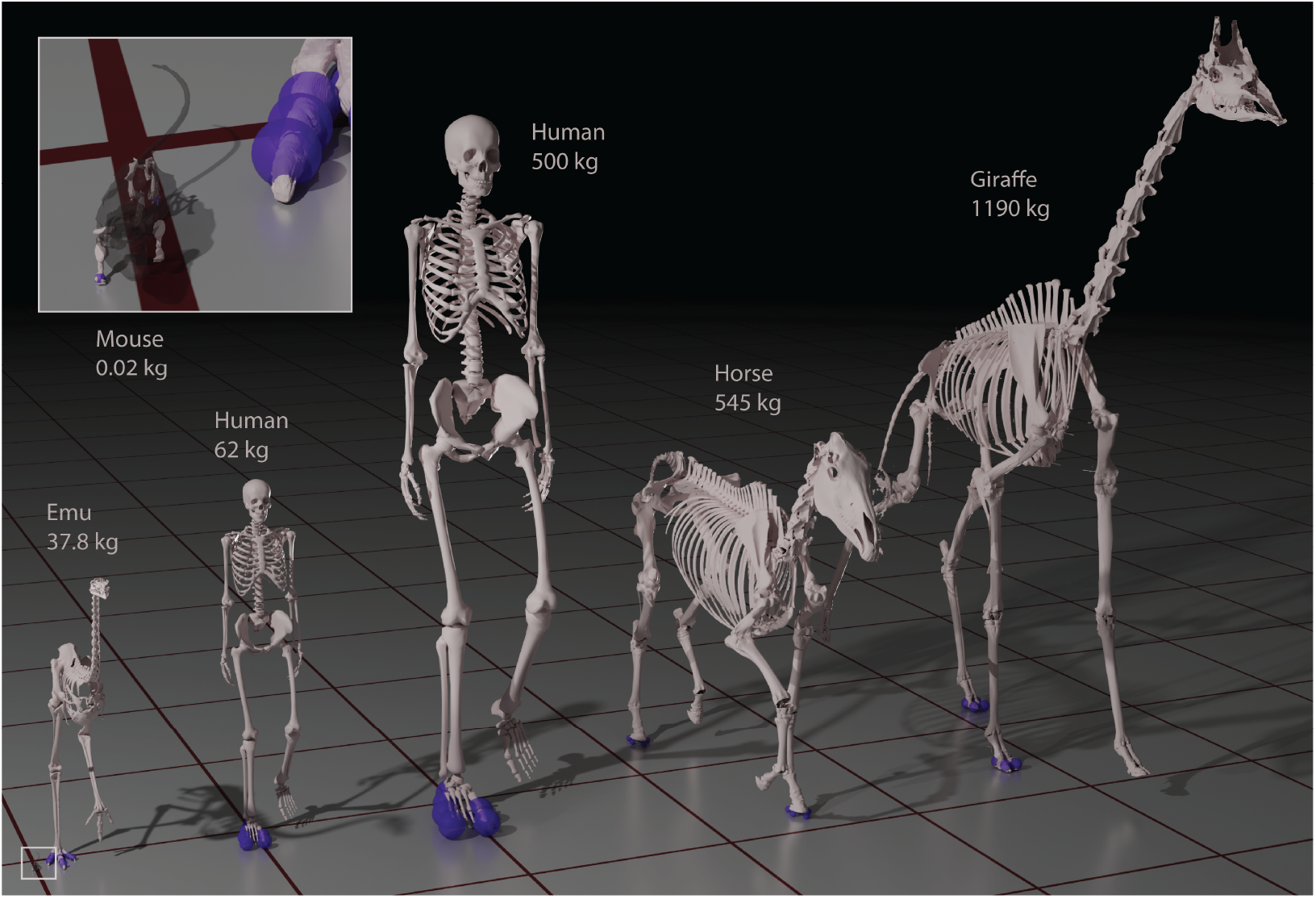
Models with different body sizes, limb counts, and numbers of contact spheres (in purple) were subjected to a simulated drop test to evaluate the effect of scaling the contact parameters. Two variants of the emu model were simulated (with 10 and 2 contact spheres), but only one is pictured. The inset depicts the mouse model, with one of the emu toes for scale. The quadrupedal models were posed so that two limbs contacted the ground (with varying number of contact spheres in total, see Table 1). The ground plane has a 1 x 1 m grid.

## 2. Derivations

Sherman et al. (2011) described an implementation of elastic contact between two non-conforming bodies, based on Hertz theory as outlined by Johnson (1985). The elastic (velocity-independent) portion of the contact force (*F*_*e*_, in N) is calculated as:

**Table 1.** Models used for the simulated drop tests, including relevant parameters that are described in the text. Two sets of simulations were performed, one with constant *α* across all models, and one with scaled *α*. Because the final equilibrium indentaion 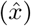 and equilibrium stiffness (*k*) were the same in both modalities, they are only reported once. ^*†*^ These models were posed so that two limbs were in contact with the ground, and the *n*_spheres_ of both feet are reported together.

| Taxon | Mass<br>(kg) | $n_{\text{spheres}}$ | $R$<br>(m) | $E^*$<br>(MPa) | $E_1^*$<br>(MPa) | base $\alpha$<br>(s m<sup>-1</sup>) | $\alpha \propto R^{-0.5}$<br>(s m<sup>-1</sup>) | final $\hat{x}$ | $k$<br>(kN m<sup>-1</sup>) |
| --- | --- | --- | --- | --- | --- | --- | --- | --- | --- |
| Human<br>*Homo sapiens* | 62 | 6 | 0.032 | 0.354 | 1.00 | 2.00 | 2.00 | 0.353 | 13.5 |
| Human<br>*Homo sapiens* | 500 | 6 | 0.0637 | 0.720 | 2.04 | 2.00 | 1.42 | 0.353 | 54.5 |
| Mouse<sup>†</sup><br>*Mus musculus* | 0.0209 | 2 | 0.002 | 0.0916 | 0.259 | 2.00 | 8.00 | 0.353 | 0.218 |
| Emu<br>*Dromaius novaehollandiae* | 37.80 | 10 | 0.015 | 0.589 | 1.66 | 2.00 | 2.92 | 0.353 | 10.5 |
| Emu<br>*Dromaius novaehollandiae* | 37.80 | 2 | 0.030 | 0.736 | 2.08 | 2.00 | 2.07 | 0.353 | 26.2 |
| Horse<sup>†</sup><br>*Equus ferus caballus* | 545 | 10 | 0.015 | 8.49 | 24.0 | 2.00 | 2.92 | 0.353 | 151 |
| Giraffe<sup>†</sup><br>*Giraffa camelopardalis* | 1190 | 8 | 0.030 | 5.79 | 16.4 | 2.00 | 2.07 | 0.353 | 207 |

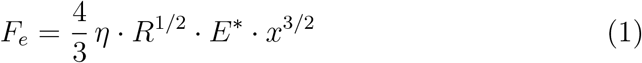

Here, *η* is 1 when using contact spheres(Sherman et al., 2011), and in general depends on the shape of the colliding objects (Johnson, 1985). Sherman et al. (2011) used the symbol *σ* instead of *η*, but in this manuscript *σ* denotes stress. *R* is a composite radius (in m); when modelling contact between a sphere and a halfspace (a plane), radius *R* is equal to the radius of the sphere (Fig. 2). *x* is the combined indentation of the two objects (in m), equal to the total intersection distance if the objects could interpenetrate. *x* thus measures how much a contact sphere deforms. *E*^*^ is the composite plane strain modulus (a measure of planar stiffness, in Nm^−2^) of the two colliding objects. By default, OpenSim assumes both the colliding objects have the same plane strain modulus 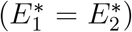, in which case both objects deform equally and the force applies halfway between both objects. Note that this implies that:

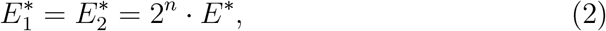

with *n* = 1 or *n* = 3/2, depending on which combining rule was used (see Appendix A for details on the combining rules, and on how to convert Young’s modulus *E* into plane strain modulus *E*^*^).

**Figure 2.**
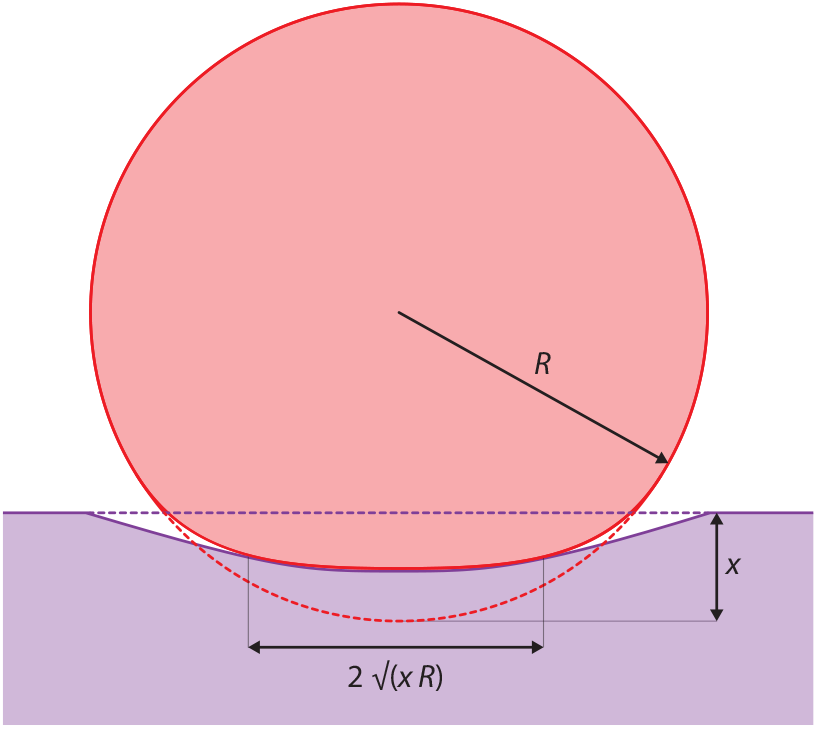
Contact interaction between a sphere with radius *R* and a halfspace. The indentation *x* is the total intersection distance between the two objects if they would not deform. The contact patch between a sphere and a halfspace is circular, with diameter 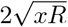.

To achieve size-invariant behaviour, it is convenient to scale *x* by *R* to acquire a dimensionless expression for relative deformation:

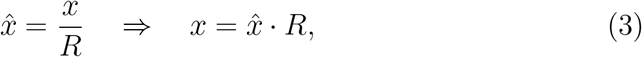

which allows us to re-express the original equation:

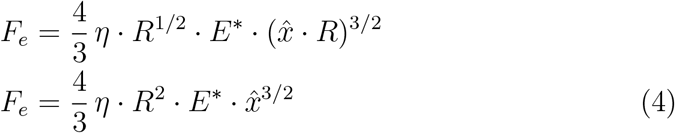

We will assume the force *F*_*e*_ represents the ground reaction force in static equilibrium, and thus scales with body mass *M*, and is equally distributed across *n*_spheres_, the investigator’s arbitrary selection of number of spheres per foot. This gives:

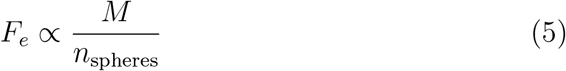

From Equation 4, the following proportionality is clear:

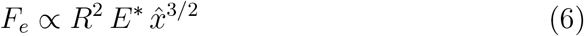

Next, we can combine this with Eq. 5 for the following proportionality:

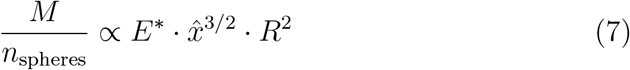

To enforce the same relative indentation irrespective of size (i.e., geometric similarity (Alexander, 2006)), we treat 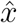 as a constant. We are left with:

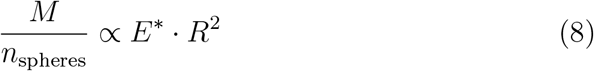

Now we can rewrite for *E*^*^, assuming constant 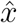:

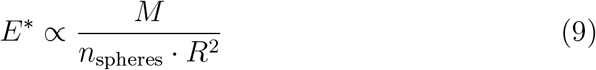

Thus, when prescribing constant relative indentation, *E*^*^ is proportional to the mass of a model, inversely proportional to the number of spheres (per limb), and inversely proportional to the square of the contact sphere radius.

### 2.1. Scaling E^*^ between models

The composite modulus *E*^*^ has the same dimensions as the plane strain moduli of the two colliding objects (Johnson, 1985; Sherman et al., 2011), and if the colliding objects have identical material properties (as assumed in OpenSim, eq. 2), then equation 9 can be used to scale the material properties. To scale *E*^*^ from model *a* to differently sized model *b*, we define the following:

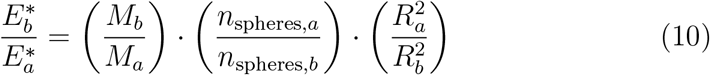

The final expression for scaling composite plane strain modulus is:

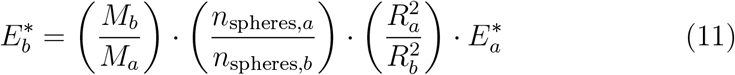

Because of the identical material assumption in OpenSim (eq. 2), this equation can also be applied to scale the individual plane strain moduli of the individual objects (i.e.,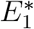), which is what the user inputs (see Appendix A).

### 2.2. Estimating a suitable E^*^ for any model

Eqs. 4 and 5 can also be combined to find a suitable *E*^*^ for a model directly, without needing to scale the parameters from a different model. This is based on the realisation that the combined forces in all the spheres need to generate a target force, which during gait would be equal to the ground reaction force. Thus, by prescribing the total force to be some multiple of bodyweight, it is possible to tailor the contact parameters for a model and a particular gait. We define:

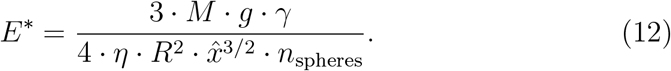

Here, *g* is the acceleration due to gravity, and *γ* is a multiplication factor for approximate peak ground reaction forces expected in the simulations (e.g., set *γ* = 2 if all the spheres together need to accommodate twice bodyweight, implying a running gait). *η* is 1 for spherical contact, and is only included for completeness.

Clearly, all the parameters on the right side are known except relative indentation 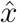. This allows the user to set 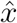 to some desired level between 0 to 1, and calculate the magnitude of *E*^*^ that would achieve this. In other words, this equation calculates the required plane strain modulus to achieve a predetermined relative indentation in the contact spheres. OpenSim users are reminded that since they supply 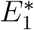, the result of eq. 12 should be multiplied by 2^*n*^ before using the result in their models (see eq. 2).

### 2.3. Constant coefficient of restitution

In simulations, damping is often included in the contact models using a Hunt Crossley formulation (Hunt and Crossley, 1975; Sherman et al., 2011; Serrancoli et al., 2019; Rodrigues Da Silva et al., 2022). In the original Hunt Crossley model, the damping force *F*_*d*_ is calculated as:

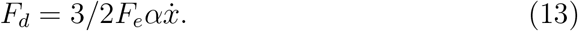

*F*_*e*_ and *F*_*d*_ are summed to acquire the total contact (normal) force. *α*, in units sm^−1^, is a parameter that is related to the coefficient of restitution *c*_*r*_ as follows:

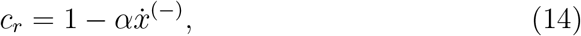

with 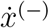 being the velocity before impact (Hunt and Crossley, 1975; Gonthier et al., 2004; Rodrigues Da Silva et al., 2022). This is an approximation that is valid for relatively low speeds and high magnitudes of *c*_*r*_ (Hunt and Crossley, 1975; Gonthier et al., 2004; Rodrigues Da Silva et al., 2022). Equation 14 shows that the (approximate) coefficient of restitution *c*_*r*_ is solely dependent on the impacting speed. However, if two differently sized animals are moving according to dynamic similarity (Alexander and Jayes, 1983; Daley and Birn-Jeffery, 2018), their speeds would be proportional to the square root of a characteristic length. In other words: 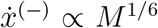, or 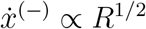. This implies that if the parameter *α* (implemented in OpenSim as “dissipation”) is kept constant for differently scaled models, the coefficients of restitution would not be identical at dynamically similar speeds. To keep the coefficient of restitution constant, 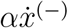 should be scale invariant. This is achieved by:

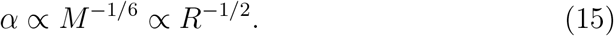

## 3. Methods

Models of a variety of different animals, encompassing a body mass of 0.02 - 1190 kg, were taken from the literature, or constructed for this study (Table 1, Fig. 1). The models were chosen to encompass a wide spread in morphologies, size, and were deliberately modelled with different *n*_spheres_. The human model from Falisse et al. (2022), scaled for Clemente et al. (2024) was provided courtesy of Christofer Clemente. The mouse model from Charles et al. (2016, 2024) was used, with a repositioned forelimb. The giraffe model was created in Blender (blender.org) using MuSkeMo (van Bijlert, 2024), and was based on a 3D surface scan of a skeleton, and a body mass measured using scales in a different specimen (euthanized for reasons unrelated to this study). All models had contact spheres on the same plane, and modifications were made using MuSkeMo if required.

The parameters for the 62 kg human model have been used succesfully in gait simulations (Falisse et al., 2022). For this study, plane strain moduli of all the other models were scaled according to equation 11 to match the 62 kg human model. Unidimensional vertical drop tests were simulated under the influence of gravity on Earth (*g*), because this experimental design isolates the dynamic behaviour of the contacts without possible interaction effects from (muscle) actuation. All models had only a single degree of freedom (vertical position), defined to be zero when the ground contact intersection commences. The final positions after 2 s were used to determine the final 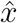, which was enough time for oscillations to have been damped out. All models were dropped from a height of 0.25*R*. To evaluate the effect of scaling *α* (the Hunt Crossley dissipation parameter), the vertical contact forces were also extracted from the simulations. These were plotted against regular time (*T*), and against non-dimensional time (Daley and Birn-Jeffery, 2018), defined as 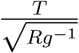.

Simulations with *α* scaled according to eq. 15 were compared to simulations where *α* = 1 across all models. Simulations were performed in Open-Sim 4.5 (Seth et al., 2018), using the Matlab (Mathworks, Natick) API. The contact forces were modelled with the “HuntCrossleyForce” contact model, which incorporates Hertz Theory and Hunt Crossley dissipative forces (Sherman et al., 2011).

## 4. Results and Discussion

### 4.1. Equilibrium behaviour

Using eq. 11 to scale the respective plane strain moduli, all the models had the same relative indentation 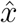 at static equilibrium (Table 1, Supplementary Figs. S1-S7). Thus, the scaling approach achieves the desired invariance to scale, morphologies, and number of legs.

All models had a relative indentation of 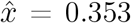, a result of scaling all models to match the 62 kg human contact parameters from Falisse et al. (2022). Note that this amount of indentation is very high — Hertz theory assumes very small deformations (Johnson, 1985; Lin et al., 2009). I am unaware of studies testing the accuracy of Hertz-theory at sphere sizes commonly used in musculoskeletal simulations, but microindentation studies suggest a maximum permissible contact radius of 0.4*R* (Lin et al., 2009). This would correspond to a maximal 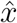 of 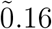. Table 1 also shows the contact stiffnesses *k* for each model (derived in Appendix B), which support the notion that the contact parameters used are relatively compliant. For example, the 62 kg human model had *k* = 13.5 kNm^−1^, which is less than a tenth of the stiffness of the heel pad in humans (Aerts et al., 1995). Despite this low stiffness, Falisse et al. (2022) achieved a very good match to empirical ground reaction forces and kinematics using these parameters in their 62 kg human model. However, D’Hondt et al. (2024) were able to improve simulated kinematics by raising *E*^*^ tenfold, and Afschrift et al. (2025) recently pointed out that the baseline contact parameters used in Falisse et al. (2022) result in supraphysiological amounts of energy dissipation in the contact model (see sec. 4.2).

It is possible to use eq. 12 to prescribe the indentation to occur at a multiple of bodyweight (e.g., 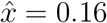 at twice bodyweight), which can keep indentations low. Furthermore, eqs. 12 and B.2 (from Appendix B) can be solved simultaneously to calculate pairs of suitable *E*^*^ and *R* for prescribed pairs of stiffness *k* and relative indentation 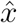. The effect would be to simultaneously prescribe stiffness and ensure indentations are within the Hertz regime. This could potentially lead to numerically undesirable stiff contact behaviour (van den Bogert et al., 1989), depending on the model and task characteristics, which may pose a pragmatic choice between parameters that abide by the small indentations assumption of Hertz contact theory or parameters that are suitable for simulations. Increasing the dissipation parameter can somewhat offset the increased simulation time of stiff contacts, but high dissipation using the Hunt Crossley model is also problematic (see sec. 4.2). High indentations can be partially avoided by enlarging the contact sphere radius, with the downside that the spheres may then become substantially larger than the impacting tissues they are meant to represent (e.g., the heel pad). Alternatively, maximizing the number of contact spheres (*n*_spheres_) for a given plane strain modulus would also serve to reduce contact indentations (eq. 12).

Hertz theory also assumes linear elastic materials (Johnson, 1985), whereas soft biological tissues tend to have non-linear elastic (“hyperelastic”) behaviour (Aerts et al., 1995; Lin et al., 2009). This suggests that the Hertz contact model is not capable of simultaneously matching measurements of tissue deformations and contact forces using measured tissue properties, especially since biological tissues generally undergo higher deformations than are permissible under the assumptions of Hertz theory. For this scenario, sphere-based contact solutions which permit large deformations and hyperelastic materials (Lin et al., 2009) could be adapted to musculoskeletal simulations.

The arguments presented above do not imply that the Hertz model must always be avoided in biological simulations. Indeed, the Hertz model’s popularity partly stems from its ability to produce realistic contact forces in a wide variety of contexts and parameter combinations (Falisse et al., 2022; Van Bijlert et al., 2024; Clemente et al., 2024; van Bijlert et al., 2024; D’Hondt et al., 2024). However, at high relative contact indentations, the model should perhaps be viewed as an abstract non-linear spring, rather than an accurate simulation of colliding spheres with prescribed material properties. The deformations can then be thought to represent the combined effects of both the material interactions, and conformational changes in the skeletal structures that in reality have more mobility than degrees of freedom in the model (e.g., models with rigid toe segments). Nevertheless, given the profound influence of substrate compliance on movement (Kerdok et al., 2002), future work could be aimed at systematically evaluating how contact parameter scaling influences movement selection in simulations.

### 4.2. Dynamic behaviour

Without scaling the dissipation parameter *α*, the dynamic behaviour of the models varied strongly with size (Fig. 3, Supplementary Figs. S1-S7). This can be most clearly observed in the mouse simulations: at *α* = 2 sm^−1^, the model oscillated around the equilibrium, and was thus underdamped with respect to the other models (Fig. 3A). After scaling *α* according to eq. 15, all models behaved in a dynamically similar manner (Fig. 3B, Supplementary Figs. S1-S7). These results demonstrate that without adjusting the dissipation parameter, models at different scales would experience proportionally different levels of energy loss during contact interactions. While this could be compensated by increased positive or negative work in the (muscle) actuators, it is often these differences in energy consumption or actuator use that are of interest in a simulation study (Afschrift et al., 2025; Van Bijlert et al., 2024). Thus, the dissipation parameter *α* should be scaled to ensure dynamically similar contact behaviour in simulations, especially when simulating models of particularly small animals.

**Figure 3.**
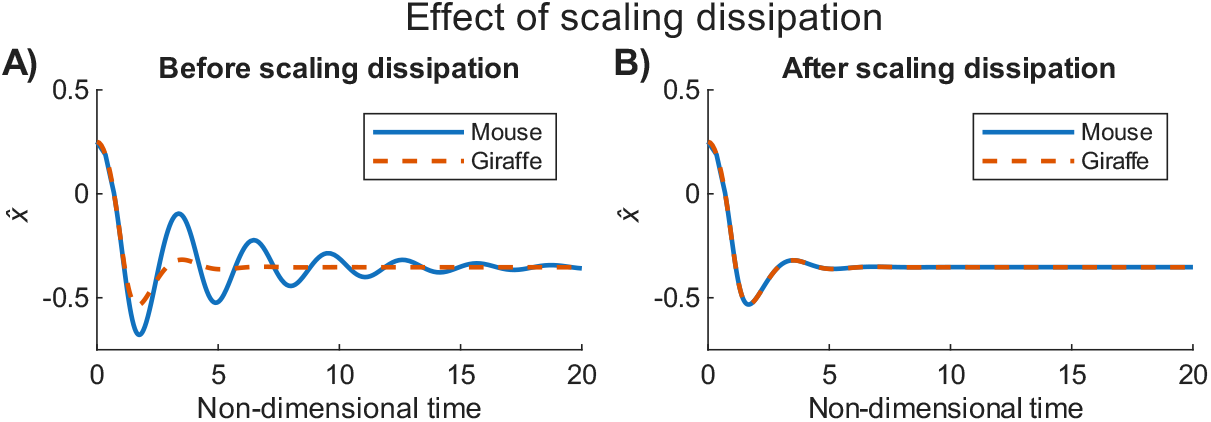
Effect of scaling contact dissipation *α*. A) Before scaling, the contact behaviour was strongly size dependent, with smaller models being progressively more underdamped. This was most extreme in the mouse model, which showed a sustained oscillation around the equilibrium indentation. B) After scaling *α*, all models showed the same dynamic behaviour, here exemplified by comparing the mouse and giraffe simulations.

The base value of dissipation used here (*α* = 2 sm^−1^) was chosen to match the human simulations by Falisse et al. (2022). This large amount of dissipa-tion violates the assumptions of the Hunt Crossley model—the Hunt Crossley model is only accurate for high coefficients of restitution (i.e., *c*_*r*_ *>* 0.8) (Hunt and Crossley, 1975; Gonthier et al., 2004; Skrinjar et al., 2018; Rodrigues Da Silva et al., 2022). The 62 kg human model had a pre-impact velocity of approximately 0.4 ms^−1^, suggesting a coefficient of restitution *c*_*r*_ of 0.2 (eq. 14). This pre-impact velocity is on the low-end for human walking (Baines et al., 2018), suggesting an even lower *c*_*r*_ in human gait simulations that used *α* = 2 sm^−1^ (Falisse et al., 2022; D’Hondt et al., 2024; Afschrift et al., 2025). As a result, contact dissipation may be inaccurately simulated when using high *α* at the scale of a human, which can have unpredictable effects on (muscle) actuation. This may have contributed to the finding by Afschrift et al. (2025) that high *α* resulted in overestimations of muscle work, and that raising *E*^*^ (while keeping *α* high and thus *c*_*r*_ low) did not fully resolve the issue.

The Hunt Crossley model’s failure to accurately model high dissipation (Gonthier et al., 2004; Skrinjar et al., 2018; Rodrigues Da Silva et al., 2022; Ding et al., 2024) is problematic for biological simulations, because dissipative contacts are necessary both for numerical stability, and to model realistic biological movements. For example, dissipation during heel-strike may account for up to one-fifth of the human metabolic cost of transport (Baines et al., 2018). Energy dissipation in the human heel pad may reach up to 65.5 % (Aerts et al., 1995), implying a coefficient of resitution that is substantially lower than 1, although the exact amount has not been measured to my knowledge. For highly dissipative contacts, alternatives or extensions to the Hunt Crossley dissipative model may be more suitable (Gonthier et al., 2004; Skrinjar et al., 2018; Rodrigues Da Silva et al., 2022; Ding et al., 2024). However, this family of contact models currently cannot account for situations where the equilibrium indentation depth changes during the contact, such as during locomotion on deformable substrates (Prescott et al., 2025) or during mastication (Watson et al., 2014), which might require a two state contact model instead.

### 4.3. Allometric implications

There is relatively little empirical data on animal limb contact mechanics to compare to, so I will compare scaling predictions with the mammalian data reported in (Michilsens et al., 2009; Chi and Louise Roth, 2010).

In static equilibrium, *F* will always be proportional to M (eq. 5). Contact stress *σ* is equal to force *F* divided by the contact area *A*. The contact area is related to *x* and *R* via:

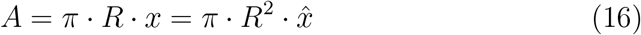

See also Fig. 2 and (Johnson, 1985). Eq. 16 holds for circular contact patches, and implies that when prescribing constant relative indentations (which only occurs with the scaling approach proposed in this manuscript), *A* scales with *M* ^2*/*3^, and thus *σ* scales with *M* ^1*/*3^. This falls in between scaling exponents reported for mammalian contact stress—*M* ^0.06^ in (Chi and Louise Roth, 2010) and *M* ^0.5^ for peak stress in Michilsens et al. (2009)—but not much work has been done on this topic. Keeping relative indentations constant implies that *E*^*^ scales with *M* ^1*/*3^ in the scaling approach proposed in this manuscript. This may seem surprising, given that the material properties of biological tissues generally do not vary with size. However, in this context the plane strain modulus *E*^*^ represents the combined effects of multiple tissues, and their relative compositions may indeed vary with size. For example, the Young’s modulus of a small sample of carnivoran footpads have been shown to scale with *M* ^0.59^ and *M* ^0.39^ (fore and hind, respectively) (Chi and Louise Roth, 2010). This supports the notion that deriving *E*^*^ based on tissue properties may have inconsistent results in musculoskeletal simulations.

*R* and *E*^*^ both scaling with *M* ^1*/*3^ implies that stiffness *k* (in Nm^−1^) scales with *M* ^2*/*3^ (see eq. B.2 in Appendix B). The literature suggests that he stiffness of carnivoran footpads scales with an exponent closer to 1 (Chi and Louise Roth, 2010). From eq. B.2, it is clear that this can only be achieved by scaling *E*^*^ ∝*M* ^2*/*3^, for a given relative indentation. In fact, relative indentation (or strain) decreases with body size in carnivorans (Chi and Louise Roth, 2010), suggesting an even higher scaling exponent for *E*^*^. Nevertheless, if we prescribe *E*^*^ ∝ *M* ^2*/*3^, once again assuming *F* ∝ *M* and *R* ∝ *M* ^1*/*3^, then by eq. 4:

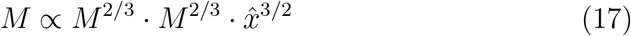

This implies that any attempt to scale the contact parameters with scale-invariant stiffness, would lead to relative indentations 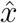 scaling with *M* ^−2*/*9^, which is correct in sign, but with a smaller magnitude than the scaling exponents found for carnivorans (Chi and Louise Roth, 2010). Furthermore, this does not achieve the goal of prescribing stiffness exactly, because constant relative indentation is invoked in eq. B.2, but not achieved as was just explained.

## 5. Conclusion

The equations and derivations presented here provide guidance for choosing scale-invariant contact parameters, following the principles of geometric and dynamic similarity. Both the plane strain modulus and dissipation parameters should be adjusted if large size-ranges are being investigated. Open-Sim’s tanh-smoothed contact model “SmoothSphereHalfSpaceForce” requires further scaling of the smoothing parameters to remain scale-invariant (see sec. Appendix C). Well-chosen contact parameters represent a tradeoff between computational efficiency and realism. Due to the assumptions of small deformations and linear elasticity, the Hertz contact model should not be expected to provide perfect correspondences with simultaneously measured biological tissue deformations and contact forces. The original Hunt Crossley dissipation model may not be adequate in simulations where high damping or other hysteresis effects are critical.

## Data availability

The OpenSim models and Matlab code to perform the simulations, and produce Figs. 3, C.4 and Supplementary Figs. S1-S7 are provided as supplementary materials. For review purposes, they are made available via: https://drive.google.com/drive/folders/1YrIs69gF4DFPLS9v45vV5p6YkQXDHVDU.

## Acknowledgements

I thank Niklas Hohmann for discussions on the initial derivations. I am grateful to Knoek van Soest for motivating me to explore the smoothed versions of the contact models, ultimately leading to recommendations presented in Appendix C. Anne Schulp and Karl Bates provided helpful comments on this manuscript. I thank Nicholas Bianco and Antoine Falisse for clarifying the contact model implementations in OpenSim. Christofer Clemente kindly shared the human models used in this manuscript, and James Charles shared extra 3D models which enabled repositioning of the mouse model. Claudia Wolschrijn of the Department of Clinical Sciences, section Anatomy and Physiology (Utrecht University) provided access to the giraffe skeleton, and Jooske IJzer and Louis van den Boom, and the staff at The Veterinary Pathology Diagnostic Centre (Utrecht University) provided the body mass measurement of a deceased giraffe.

## Appendix A Plane strain modulus

For an object with Young’s modulus *E*_1_ and Poisson’s ratio ν_1_, its plane strain modulus 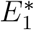 is computed as:

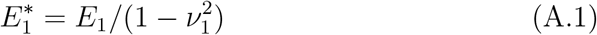

The composite plane strain modulus *E*^*^ can be constructed from the plane strain moduli of the two colliding objects. *E*^*^ is constructed using either (Johnson, 1985):

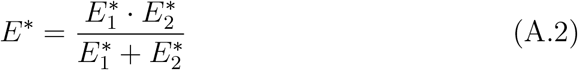

Alternatively, using the combining rule derived by Sherman et al. (2011), which is implemented in Simbody and OpenSim:

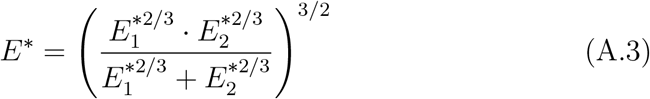

By default, OpenSim assumes that both the colliding objects have the same plane strain modulus 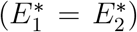, in which case both objects deform equally and the force applies halfway between both objects. Note that this implies that:

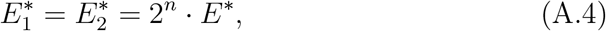

with *n* = 1 when using eq. A.2 and *n* = 3/2 when using eq. A.3. This has consequences if the intended goal is to model the interactions using measured or estimated material properties, instead of the functional approach proposed in the main manuscript. For example, if the intention is to model the interaction between an animal’s foot pad (i.e., a soft object with measured Young’s modulus *E*_1_, *m*), and a force plate or concrete (an object assumed to have effectively infinite stiffness), then in the limit for 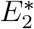 to infinity in eqs. A.2 and A.3:

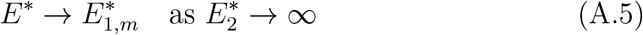

Because in OpenSim, the input is 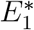 and not *E*^*^, combining eqs. A.1, 2, and A.5, OpenSim users should input 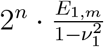 as the plane strain modulus when starting from a measured Young’s modulus, under the assumption that the other surface has an effectively infinite Young’s modulus. Note that the force would still be applied halfway between the two colliding objects.

## Appendix B Contact force stiffness

Equation 1 can be viewed as a non-linear spring. Stiffness *k* then follows as 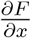:

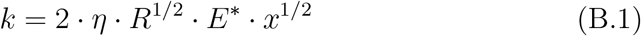

Combining this with eq. 3, we acquire:

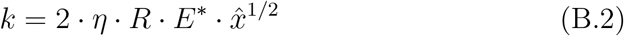

This shows that effective stiffness is linear in both *R* and *E*^*^, at any given relative indentation. It is possible to solve eqs. 12 and B.2 as simultaneous equations, and use this to estimate suitable values for both *E*^*^ and *R*. This can be achieved by prescribing a desired stiffness *k* (in Nm^−1^ and a (maximum) relative indendation 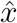, and then solving for the unknown *E*^*^ and *R*.

## Appendix C SmoothSphereHalfSpaceForce in OpenSim

OpenSim includes “SmoothSphereHalfSpaceForce”, which is a smoothed version of the standard HuntCrossleyForce, developed specifically for increased performance in gradient-based optimisations (Serrancoli et al., 2019; Dembia et al., 2020). The elastic (Hertz) portion of the contact model is smoothed in two ways: 1) by addition of term *cf* that ensures a low force even when there is no contact (thus preventing null-derivatives), and 2) by approximating the binary conditions of no contact/contact with a tanh function based on the contact indentation (preventing infinite derivatives). The entire model is also smoothed at the velocity level with a second tanh function to prevent negative contact forces. The smoothing parameters are not dimensionless, and thus their effects are sensitive to the scale of the contact parameters, unless the smoothing parameters are also scaled. Fig. C.4 shows that inappropriate combinations of parameters can lead to high contact forces well-before the actual contact is initiated (signified by a the red shaded area in panel A). This can be interpreted as the contacts repelling the ground without contact, or alternatively as modelling larger contact spheres than intended. This issue occurs when the *cf* term is relatively high, combined with too smooth a tanh-transition, which together allow the constant contact force to bleed through before contact is actually initiated.

**Figure C.4.**
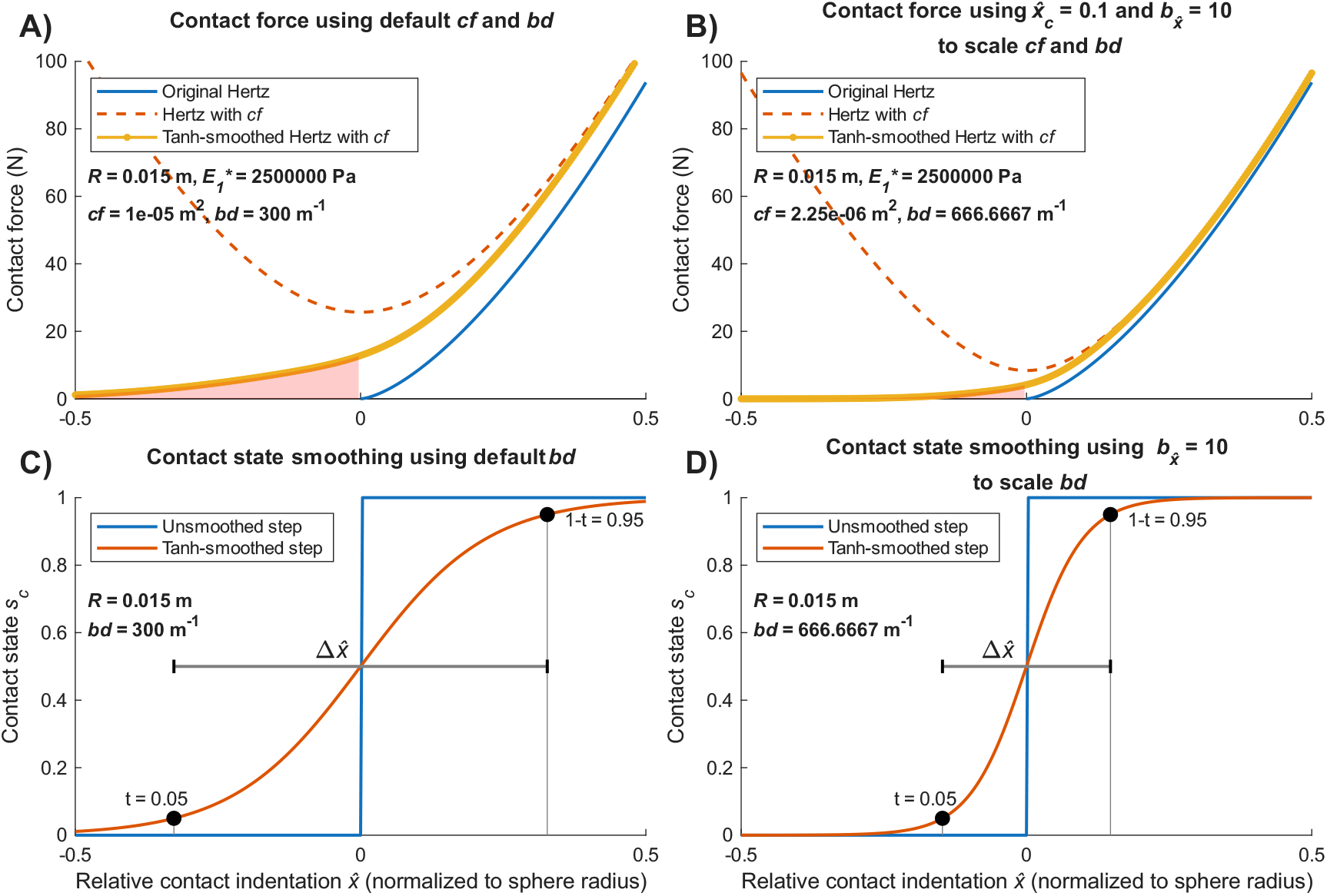
The smoothing parameters in OpenSim’s “SmoothSphereHalfSpaceForce” are not dimensionless, and can cause issues when using relatively small contact spheres (here, *R* = 0.015 m). A) Using default values for the *cf* and *bd* parameters, the smooth contact model (yellow dotted line) produces substantial amounts of force before contact is initiated, signified by the red shaded area. The *cf* parameter, which effectively adds a constant contact indentation, resulting in the Hertz force with *cf* (dashed orange line) being substantially higher than the regular Hertz force (blue line). Force builds up before contact initiation due to smoothing via *bd* which results in contact state buildup before physical contact (see panel C). B) After scaling, the smoothed force is much closer to the unsmoothed Hertz force, because reduced *cf* reduces the constant contact force, and increased *bd* reduces the transition width (see panel D). C) In the unsmoothed model, contact is a binary state, signified by a step function. This is smoothly approximated by a tanh function, which means contact is initiated before the sphere intersects the halfspace. The default value of *bd* encodes an absolute transition distance, which means that in small contact spheres, the smoothing has a much stronger effect. D) After scaling *bd*, the contact transition occurs over a much smaller fraction of the sphere radius. This prevents large forces bleeding through before contact is initiated.

An interactive way to check whether this issue is present in a model is to load it into OpenSim Creator (Kewley et al., 2024), and to observe whether the real-time contact force visualisations build up substantially before the contact spheres intersect the contact halfspace. A Matlab script that parametrically generates Fig. C.4 is provided in the supplementary materials, which enables plotting the effects of different scaling choices. In the next sections, I will present some derivations and physical interpretations of the smoothing parameters. I will also provide some practical recommendations on how to scale them, based on published models where the smoothing parameters apparently caused no issues.

## Appendix C.1 Constant force due to cf

Inclusion of the parameter *cf* results in a rewritten version of eq. 1 (Serrancoli et al., 2019):

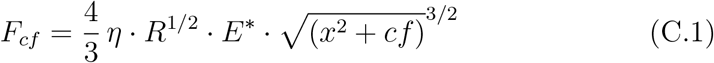

While the parameter *cf* is named “constant_contact_force” in OpenSim, from its position in eq. C.1 it is perhaps more accurately described as a “constant contact indentation squared”, with units m^2^. This is an important distinction, because the indentation due to *cf* can result in drastically different force magnitudes depending on the contact radius (eq. C.1). Open-Sim’s default value of 10^−5^ m^2^ corresponds to a constant contact indentation of *x* = 0.0032 m, and it is advisable to scale the magnitude of *cf* with *R*. For instance, the mouse model simulated in this manuscript has *R* = 0.002 (m). The default *cf* would result in an indentation of more than 1.5 times the contact radius, causing the model to float in the air during steady-state. This would be obvious in such a small model, but is much more insidious in a medium sized model. For instance, the emu model used *R* = 0.015 m, and this issue can only be diagnosed by plotting the force indentation curve (fig. C.4). One scaling approach would be to choose *cf* based on a desired level of constant relative indentation 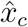 (which is a fraction of *R*, and thus dimensionless):

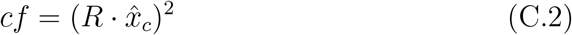

Lower 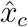 then results in less constant indentation, and vice versa. Using the Falisse et al. (2022) human model’s *R* = 0.032 m as a reference value, a possible starting point for 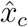 could be 0.0988, or simply 0.1, signifying a constant contact deformation of 10% of the contact radius. Fig. C.4B demonstrates the effect of this scaling. If the recommendation of 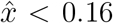 from the main manuscript is followed, 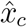 should be adjusted accordingly (e.g., 0.05).

## Appendix C.2 Smoothing with tanh functions

The Hertz equation (eq. 1) is only defined when the objects are actually in contact. Simulators can enact this by only computing the force if *x* is positive (negative *x* thus represents no contact). Such an “if statement” can be thought of as a binary step—when a contact sphere goes from zero indentation to a finite level of indentation, there is a binary step in the contact state from 0 to 1 (C.4C). The binary step (between 0 and 1, representing “no” and “some” contact, respectively) can be smoothly approximated by introducing a smoothed contact state *s*_*c*_, which depends on *x*. Then, the smoothed Hertz force *F*_*s*_ can be computed as follows:

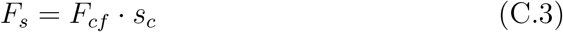

In the unsmoothed model, contact is binary, and thus no forces are produced when there are no intersections. In the smoothed model, the contact state *s*_*c*_ is continuous between 0 and 1, as a function of indentation *x*. In SmoothSphereHalfSpaceForce (Dembia et al., 2020; Serrancoli et al., 2019), the smoothing is achieved with a tanh function (Fig. C.4C, plotted as a function of 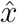):

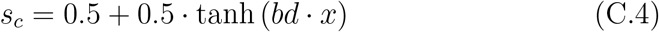

The parameter *bd*, called “hertz_smoothing” in OpenSim, decides how smooth the transition is between 0 and 1. By the nature of this function, *s*_*c*_ tends asymptotically to 0 for negative *x*, asymptotically to 1 for positive *x*, and is exactly 0.5 when *x* = 0. In other words, the tanh-approximation results in the contact state building up before the sphere actually intersects the halfspace. Because *F*_*cf*_ is non-zero before the spheres intersect (due to the constant force due to *cf*), the smoothed contact force *F*_*s*_ will also be non-zero before the spheres intersect (eq. C.3). While this behaviour is intentional (to improve gradient based optimizations (Serrancoli et al., 2019)), it is not scale invariant: eq. C.4 implies *bd* has units m^−1^, the reciprocal of the length scale, because *s*_*c*_ itself is dimensionless. Thus, the smoothing enacted by *bd* is not dimensionless, the transition from “no contact” to “contact” will always occur over the same absolute distance, whereas the user is likely to tailor *R* to the scale of the model.

The OpenSim default value of parameter *bd* (300 m^−1^) will have relatively little smoothing effect for large *R* (e.g., *R >* 0.06 m), but small values of *R* (e.g., *R <* 0.02 m) are more problematic, especially if *cf* was not scaled (compare the shaded areas in Fig. C.4A and B). Based on its placement in eq. C.4, it is clear that *bd* should be scaled according to *bd* ∝*R*^−1^. This can be used to scale *bd* between different models. To specify an arbitrarily steep transition:

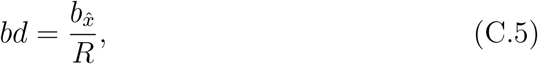

where 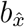 is a dimensionless smoothing parameter which controls the relative indentation 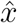 over which any given fraction of the *s*_*c*_ step occurs (derived below). The Falisse et al. (2022) model does not have the smoothing issue, and used *bd* = 300 m^−1^, with *R* = 0.032 m. This implies that a suitable starting value for 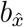 could be 300 · 0.032 = 9.6, or simply 10. Setting 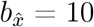 would imply that the step function increases from 5% to 95% when 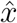 varies from −0.15 to 0.15 *R* (see eq. C.8 below, and Fig. C.4C and D).

Eq. C.4 can be rewritten to estimate a suitable 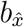 with more fine-grained control, which allows specifying a percentage of the step that should be approximated by a given fraction of the radius. To do this, we introduce a variable *t* that enables us to define a threshold on our our step function. For instance, if we desire our step function to reach 95% for a given relative indentation 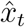, we choose *t* = 0.05, and set our threshold as follows *s*_*c*_ = 1 − *t* (Fig. C.4C and D). We can insert this into eq. C.4, and solve for 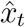, which gives:

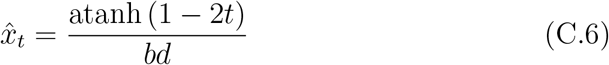

Because eq. C.4 is symmetric about the point (0, 0.5), we can use the above to define a width in 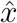 over which the transition between *s*_*c*_ = *t* and *s*_*c*_ = 1 − *t* occurs:

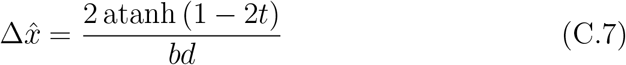

For example, if 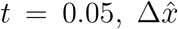 defines the negative to positive indentation which encompasses 90% of the total step function (because it is symmetric about 0.5, Fig. C.4C and D). Rewriting the above equation to solve for *bd*, and once again using eq. 3, we acquire:

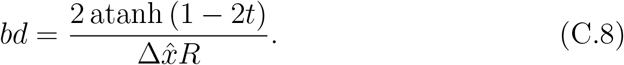

This shows that 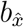 from eq. C.5 can be parametrised as 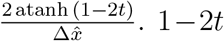 can be interpreted as the percentage of the step that is traversed over 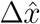, the relative indentation over which it occurs. In the limit for 1 − 2*t* to 1 and 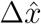 to 0, the step function is no longer an approximation, but also no longer smooth. The supplementary files contain a Matlab script that allows plotting the effects of different combinations of 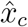 and 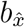.

In SmoothSphereHalfSpaceForce, the total normal force is subjected to a second layer of smoothing, with 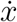 as the input variable (Serrancoli et al., 2019; Dembia et al., 2020). This is to ensure that the dissipative (Hunt Crossley) force can never cause negative contact forces (sticking). This is achieved by centering the tanh at the speed where *F*_*d*_ = - *F*_*e*_, which occurs at 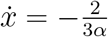 (see eq. 13), and introducing a second smoothing parameter *bv*, called “hunt_crossley_smoothing” (default value = 50 sm^−1^). In equation form:

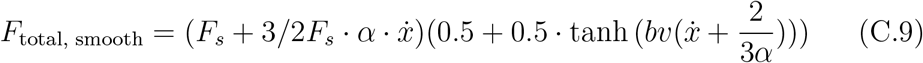

The term between the first brackets is the smoothed equivalent of *F*_*e*_ +*F*_*d*_, and the second term is the velocity smoothing term. The full derivation is left as an exercise to the reader, because by analogy with eqs. C.5 and C.8, we can see that *bv* has units sm^−1^, scales according to *bv* ∝*R*^−1*/*2^, and can be customised by the user as follows:

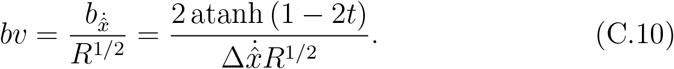

Again referencing the Falisse et al. (2022) model, a starting point for 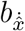. could be 50 · 0.032^1*/*2^ = 8.94, or simply 9.

Scaling *bv* may have very little effect in practice, unless the movement that is being simulated has very high negative 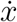 (e.g., a high takeoff velocity).

## Footnotes

1 Hertz derived what is now known as Hertz theory of elastic contact at the age of 23 during his Christmas break in 1880, essentially starting the field of contact mechanics (Johnson, 1985). He is better-known for conducting experiments demonstrating the existence of electromagnetic waves (Hertz, 1893), which is honoured by the eponymous SI-unit Hz

